# Microbial Community multi-omic analysis of marsh sediment post crustacean shell compost enrichment: pathogen emergence and community response

**DOI:** 10.64898/2026.03.06.710096

**Authors:** Colleen E. Yancey, Kyle D. Brumfield, Laurence Ettwiller, Rita R. Colwell

## Abstract

Changes in nutrient availability can rapidly alter microbial processes in natural environments, with implications in biogeochemical cycling and pathogen emergence. Short-term, functional responses of microbial communities to nutrient amendment in coastal communities remain poorly understood, particularly in temperate environments. A 48-hour microcosm pulse experiment was completed in which paired metagenomic and metatranscriptomic sequencing were employed to examine how the decomposition of chitin rich substrates, namely crab and lobster shell compost, alters salt marsh microbiome structure and function. Within 48 hours of amendment, pronounced shifts in community metabolism were observed, including increased chitin degradation and utilization, stress-response, and sporulation. These responses coincided with marked decreases in genes associated with key biogeochemical processes, including carbon fixation, sulfur oxidation and reduction, and other metabolic pathways. Shell compost addition also enriched putative pathogens and virulence-associated genes, accompanied by modest transcriptional activation, notably aerolysin A *(aerA)*, which encodes the pore-forming exotoxin aerolysin. These results demonstrate temperate salt marsh sediment microbiomes can undergo shifts in community composition and function that is associated with chitin-rich nutrient perturbation. The sensitivity of temperate coastal systems to organic matter input and the potential for ecological and public-health relevant outcomes are underscored, notably given that chitin is among the most abundant and readily available bionutrients in aquatic ecosystems globally.

## Introduction

Coastal environments, including salt marshes and estuaries, are a source of ecosystem services, including biogeochemical cycling, fishery nurseries, other aquaculture, and buffer zones during extreme precipitation events [1, 2]. These coastal systems harbor rich and largely uncharacterized microbial diversity, significant in biogeochemical cycling of carbon, nitrogen, and sulfur [3–5]. Of note is the importance of salt marshes in carbon turnover and sequestration, largely driven by diverse microbial communities inhabiting these environments [5, 6]. However, while these microbial communities provide a variety of services that benefit the health of the ecosystem and that of society, they also may harbor potentially harmful bacteria and toxic algae [7–10].

The occurrence of potentially harmful microorganisms in coastal systems is increasing, both in range and duration, as a result of a warming climate [11–15]. The incidence of *Vibrio* species, including those responsible for cholera and vibriosis, is intensifying in aquatic ecosystems worldwide, including the Chesapeake Bay, Maryland, USA [16], Florida Gulf Coast [17], and northern latitudes [18–20]. Other taxa of public health significance include *Aeromonas* spp., responsible both for major economic loss in aquaculture and life-threatening infection in immunocompromised individuals [21–23]. While these species are autochthonous in low abundance in aquatic ecosystems, their emergence and proliferation can be exacerbated during hurricanes and extreme flooding events [7, 17, 24, 25].

The second most abundant natural polymer globally is chitin, a bioavailable polysaccharide found in the exoskeleton of crustaceans (copepods, shrimp, crab, etc.), insects, and fungal species. Chitin plays an important role in both carbon and nitrogen biogeochemical cycling. Comprised of β-(1→4) linked N-acetyl-D-glucosamine, chitin does not accumulate in natural systems, likely due to efficient degradation and turnover [26, 27]. It serves as a nutrient rich substrate for a variety of microbial taxa [26, 28], including potentially pathogenic bacteria colonizing shellfish and other chitin rich surfaces in the natural environment [29–32]. Although salt marshes provide critical buffer zones during flooding, they are vulnerable to nutrient loading, particularly with organic matter accumulating during extreme weather events such as hurricanes [33]. During these events, microbial growth is accelerated, and opportunistic and pathogenic taxa can emerge [17]. The increased frequency of extreme events requires improved understanding of nutrient loading, and how it relates to microbial community composition and functional alterations

The Great Salt Marsh, spanning the North Shore of Massachusetts and the New Hampshire Seacoast, comprises more than 20,000 acres and forms a vital ecosystem of the region. Like other northern salt marshes, it experiences strong seasonal variation, with markedly reduced productivity during the winter months, unlike southern marshes which maintain higher productivity year-round [34]. Additionally, the sediment within this system contains high levels of organic carbon compared to other salt marshes of the Eastern United States [35], suggesting efficient accumulation and accretion. However, long-term carbon storage in these sediments is declining, as eutrophication reduces root biomass and organic matter accumulation, potentially leading to marsh elevation loss [36]. The Great Marsh offers a unique site for studying microbial population dynamics responding to temperate system nutrient loading and declining carbon sequestration.

The objective of this study was to investigate the effects of nutrient amendment on salt marsh soil microbial community structure and function by performing microcosm pulse experiments and multi-omic analysis. Specifically, the transcriptional response of the entire microbial community fed a chitin rich substrate, composed of crab and lobster shell compost (subsequently referred to as shell-compost), was investigated in a data-driven, hypothesis agnostic manner. The design rationale was to determine rapid shifts in community composition in response to nutrient loading. Results highlight the dynamic response of the microbial community within 48 hours, characterized by shifts to chitin degradation and stress responses, in addition to increased abundance of specific pathogenic taxa. The results provide insights into the microbial response to different nutrients with high resolution and are informative for future management practices.

## Materials and Methods

### Site description and sample collection

Soil samples were collected from the Great Marsh, Ipswich MA, USA (42°40’20.5”N 70°48’31.8”W), which spans the North Shore of Massachusetts. The area is considered an open marsh that is drained by the Labor-in-Vain Creek and experiences flooding during high tide and drainage during low tide. The site is largely dominated by high marsh (elevation greater than 1.3m) and experiences semidiurnal tides with an average mean range of 2.5-2.9 m [37]. The marsh is dominated by saltmeadow cordgrass, *Sporobolus pumilus* [35, 38], and the sediment contains high levels of organic carbon, compared to other salt marshes of the eastern United States [35].

Sampling was done on October 2, 2023, at 7:15 am during low tide. At the time of sample collection, the air and soil temperatures were 16°C and 15°C, respectively. Soil pH, determined using VWR pH-Test strips (BDH35309.606), was approximately 6. Soil samples were collected from the mud flats, at 0-10cm depth, adjacent to *S. pumilus* growth, a habitat suitable for chitin-rich organisms, namely crustaceans and mollusks [39]. Collected in bulk, the soil samples were stored in sterile high-density polypropylene (HDPE) bottles at ambient temperature for transport to the laboratory. Samples were immediately processed upon arrival, approximately 20 minutes post sample collection.

### Microcosm experiments

The collected soil samples were homogenized and passed through a 2.36 mm sieve to remove larger debris. Two microcosms were prepared, each containing 600g of homogenized soil in separate, sterile, glass containers. Both microcosms were maintained at room temperature (25°C) for the duration of the study. One microcosm served as a control, containing only homogenized marsh soil. The second was amended with 10% w/w sterile (autoclaved), ground, commercially available crab and lobster shell compost. Shell compost was chosen over purified or synthetically derived chitin to more accurately reflect the types of substrates occurring in natural environments. It is also worth noting that purified/chemically derived chitin is less preferred for degradation by microorganisms [40, 41]. The microcosms were mixed using sterile spatulas and destructively sampled at predetermined time points (0, 1, 4, 8, 24, and 48 hours), by collecting two-gram aliquots in duplicate. Extracted samples were flash frozen using an ethanol/dry ice slurry and stored at –80°C until further analysis.

### Preparation of Nucleic Acid

A total of 40 samples, including control and treatment microcosms recovered from six time points in technical duplicates, were selected for shotgun metagenomic sequencing. Total DNA was prepared from 0.25 g of soil per replicate using DNeasy PowerSoil Pro Kit (Qiagen, Hilden, Germany, 47014). OneStep PCR Inhibitor Removal Kit (Zymo Research Corp, Irvine, CA, USA, D6030) was used to treat and concentrate the DNA preparations. Double-stranded DNA concentrations and purity ratios were determined using NanoDrop One Microvolume UV Spectrophotometer (ThermoScientific, Waltham, MA, USA).

RNA for metatranscriptomic sequencing and analysis was extracted and prepared as described. RNeasy® PowerSoil® Total RNA kits (Qiagen, Hilden, Germany, 12866-25) were used to extract total RNA from 2 grams of soil for each replicate, following manufacturer’s instructions, with the following modification. The lysis/bead bashing step was increased to 20 minutes. During isopropanol precipitation, samples were incubated at –20°C for 45 minutes. To remove residual polyphenolic or humic acids that would interfere with downstream library preparation, OneStep PCR Inhibitor Removal Kit (Zymo Research Corp, Irvine, CA, USA, D6030) was used to treat and concentrate purified RNA following manufacturer’s instructions. RNA concentration and purity ratios were determined using NanoDrop One (ThermoScientific, Waltham, MA, USA). Samples were stored at –80°C until continuation of library preparation.

### Library Preparation and Sequencing

For each metagenomic sample preparation, 50 ng of purified DNA was sheared for 200 bp inserts using the Covaris S2 Focused Ultrasonicator (Covaris, Woburn, MA, USA). Sequencing libraries were prepared using NEBNext Ultra™ II DNA Library Prep Kit for Illumina (New England Biolabs, Ipswich, MA, USA, E7645). NEBNext Multiplex Oligos for Illumina kit (New England Biolabs, Ipswich, MA, USA, E6444) were used for indexing and PCR amplification (5 cycles). Library concentration, size, and quality were assessed using D1000 High Sensitivity Screen Tape with the TapeStation 4200 (Agilent, Lexington, MA, USA, 5067-5585). DNA libraries were sequenced at the New England Biolabs Sequencing Core, using NovaSeq S2/SP System (Illumina, San Diego, CA, USA) with 100 bp paired end reads. Shotgun metagenomics was done targeting >60M paired reads.

For each metatranscriptomic sample, ∼800 ng of total RNA was used for library preparation. Ribosomal depletion was completed using NEBNext® rRNA Depletion Kit for Bacteria (New England Biolabs, Ipswich, MA, USA, E7850), following the manufacturer instructions. Libraries with ∼200 bp inserts were prepared with NEBNext® Ultra II Directional RNA Library Prep Kit (New England Biolabs, Ipswich, MA, USA, E7765). NEBNext® Multiplex Oligos for Illumina® (New England Biolabs, Ipswich, MA, USA) were used for indexing and PCR amplification (6 cycles). Library concentration, size, and quality were assessed using a TapeStation 4200 with a D1000 High Sensitivity Screen Tape (Lexington, MA, USA). Shotgun metatranscriptomics was completed targeting >100M paired reads.

### Preprocessing of sequencing reads

Quality trimming and adaptor removal from paired read libraries was completed with trimgalore v.0.6.10 [42] under default settings. Trimmed read files were inspected using FastQC v.0.12.1 to assess sequence quality.

### Whole community microbiome profiling

Metagenomic and metatranscriptomic reads were profiled using Kraken2 v.2.1.3 [43] under default settings with the Standard Collection database built for 100-mers (accessed October 2024). Taxonomic profiles were visualized using Phylosmith v1.0.6 [44] and presented as relative sequencing read abundance.

Depth of the sequencing read libraries was normalized to the sample with minimum number of simulated reads using the ‘rarefy_even_depth’ function of phyloseq v1.46 [45]. Alpha diversity metrics were obtained using the phyloseq ‘estimate_richness’ function. The observed (number of taxa), Shannon (entropy) [46], and inverse Simpson’s diversity index (richness under uniform evenness) [47] were visualized using ‘plot_richness’ of phyloseq. In addition, raw count data were transformed into relative abundances, expressed as counts per million (CPM), and used for principal coordinate analysis (PCoA) employing Bray-Curtis [48, 49] distance measure through phyloseq’s ‘ordinate’ function. Differential abundance analysis was performed using DESeq2 v1.42.1 [50]. A DESeqDataSet object was generated from normalized bacterial abundance (CPM) using the ‘phyloseq_to_deseq2’ function. Generalized linear models were fitted for various time points using control samples as reference by calculating the ratio of their likelihoods. Resulting P-values were adjusted using the Benjamini-Hochberg procedure [51] (FDR ≤ 0.01). Taxa were considered significant if they exhibited differential abundance at late timepoints (24 and 48 hr) relative to early timepoints (1 and 4 hr). Results are reported as log_2_ fold changes.

### Transcriptional Profiling in response to treatment

To assess the effects of shell compost addition on community dynamics, a variety of gene families containing functional domains of interest were analyzed through the metaGPA pipeline. Assembled transcripts were annotated using the Protein Family Database, PFAM [52], v 35.0, and Hidden Markov modelling (HMM) to complete annotation via hmmer v 3.3.2 under default settings. This analytical pipeline is described in greater detail elsewhere [53]. Domain annotations with a p-value of at least 1e-05 were considered significantly associated.

Sequences containing domains of interest with a p-value of 0 were extracted and Normalized Transcriptional Changes calculated. First, to correct for changes in community composition over time, RNA relative abundance was normalized to DNA relative abundance using the following equation:

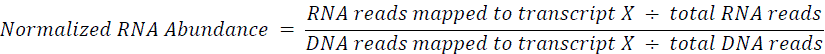

Next, the ratio between Normalized RNA Abundance in Treatment and Control for each transcript at each corresponding time point was calculated and log_2_ transformed. The following equation was used to calculate Normalized Transcriptional Changes between treatment and control samples:

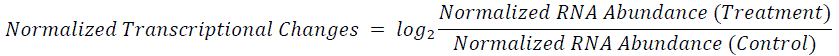

### Data Visualization

Statistical analysis and figure rendering were completed using R and R Studio v.4.4.2 via the tidyverse collection.

### Gene profiling for antimicrobial resistance and virulence determinants

Presence and relative expression of antimicrobial resistance genes and virulence determinants were profiled and quantified using shortBRED. Briefly, custom cluster databases were constructed using coding sequences from the NCBI Pathogen Detection Reference Gene Catalog [54] and Virulence Factor Database [55] and the ‘-shortBRED-Identify’ function. Sample reads were mapped against the databases using ‘-ShortBRED-Identify, relatively quantified, and reported as reads per kilobase of transcript per million reads mapped (RPKM). Sequences that were present in at least one sample with an RPKM value of at least one were retained for further analysis.

## Results

Salt marsh soil was collected from the Great Marsh, homogenized, and either amended with shell compost (treatment microcosms) or left unamended (control microcosms). Duplicate samples were collected at 0, 1, 4, 8, 24, and 48 hours after the amendment step to assess effects of shell compost addition on microbial community dynamics. DNA and RNA were extracted from all samples and respectively subjected to metagenomic and metatranscriptomic sequencing to assess changes in community composition and transcriptional activity.

### Shell Compost Addition Alters Species Diversity and Community Composition

Microcosm microbial communities exhibited dynamic shifts in taxonomic composition at later time points for samples treated with shell compost compared to untreated samples. Measurements of alpha diversity indicated a decrease in species diversity for samples collected 24 and 48 hours after shell compost addition (Fig. 1A). Beta diversity, measured by Bray-Curtis dissimilatory, indicated similar community composition of samples collected before 24 hours regardless of treatment, while samples collected at 24 and 48 hours were distinct relative to sampling time and treatment (Fig. 1B). Shifts in bacterial community composition 24-48 hours after shell compost addition were largely driven by a relative increase in Gammaproteobacteria and Bacilli, consistent with previous analyses [53]. An increase in Epsilonproteobacteria was also observed at 48 hours in both treatment and untreated control samples (Fig. 1C). Several bacterial genera represented this compositional shift, including notable increase in *Vibrio*, *Shewanella*, *Aeromonas*, and *Bacillus spp.*, with *Acinetobacter* spp. also increasing at 48 hours (Fig 1D,E). In contrast to treated samples, untreated samples remained relatively stable in community composition from time 0 to 48 hours (Fig. 1).

**Figure 1:**
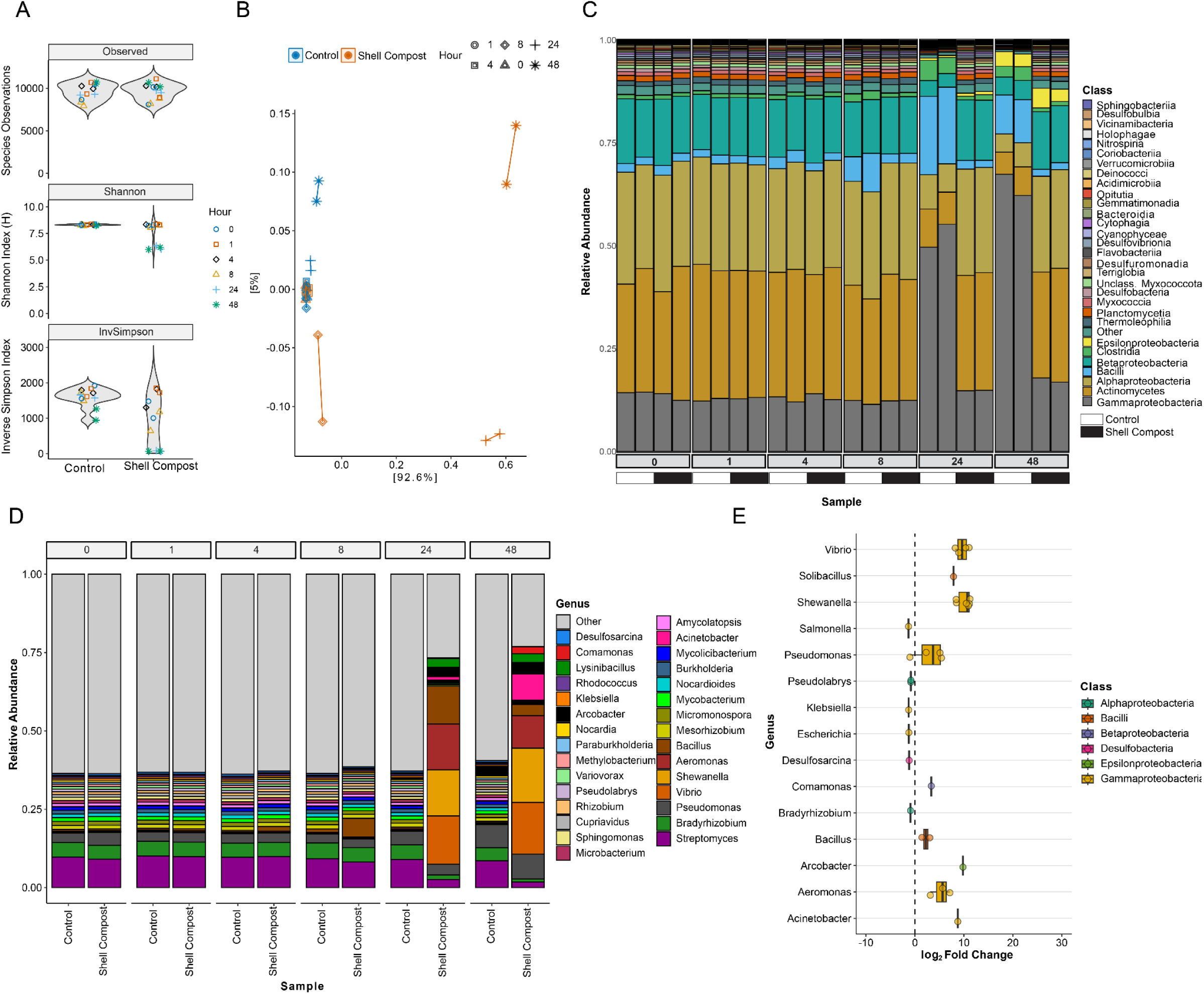
Microbial community composition. (A) Violin plots showing alpha diversity between microbiome taxonomic profiling sets. (B) Principal coordinate analysis of microbial communities. (C) Stacked bar plot showing relative sequencing read abundance of bacterial classes for each replicate sequenced. Black and white boxes are used to denote shell compost treated and control samples, respectively. (D) Stacked bar plot showing averaged relative sequencing read abundance for bacterial genera. The top 30 most abundant genera are shown, while the rest are denoted as “Other” (colored gray). (E) Boxplot showing likelihood ratios of genera among most abundant genera.

### Compost Addition Alters the Soil Microbiome Transcript Pool

Metatranscriptomic read profiling exhibited distinct patterns, representing differences in microbial community composition and transcriptional response. For example, metagenomic and metatranscriptomic read profiling did not exhibit a 1:1 relationship for several taxa, including *Vibrio* and *Aeromonas* spp., which increased in abundance but represented only a small portion of total transcripts. Transcriptional alpha diversity was similar among time points regardless of treatment, although indices were lower compared to metagenomic read profiling (Fig. 1A, 2A). Beta diversity revealed more similar transcript pools for earlier timepoints (0-8 hours), regardless of treatment, with distinct transcript pools at 24-48 hours for both control and treated samples (Fig. 2B). Compared to metagenomic read profiling, shifts in the transcript pool were less dynamic, with Gammaproteobacteria, Bacilli, and Clostridia comprising >40% of the profiled transcripts across samples. The genera *Pseudomonas* and *Bacillus* comprised ca. 25% of the transcript pool, regardless of sampling type (Fig. 1C, 2C). However, at 24 and 48 hours after shell compost addition, *Acinetobacter, Lysinibacillus, Clostridium, Aeromonas*, and *Solibacillus* spp. notably increased in relative transcript abundance. *Acrobacter* spp. and *Commamonas* spp. transcripts increased in control and shell compost treated samples respectively at 48 hours (Fig. 2D,E).

**Figure 2:**
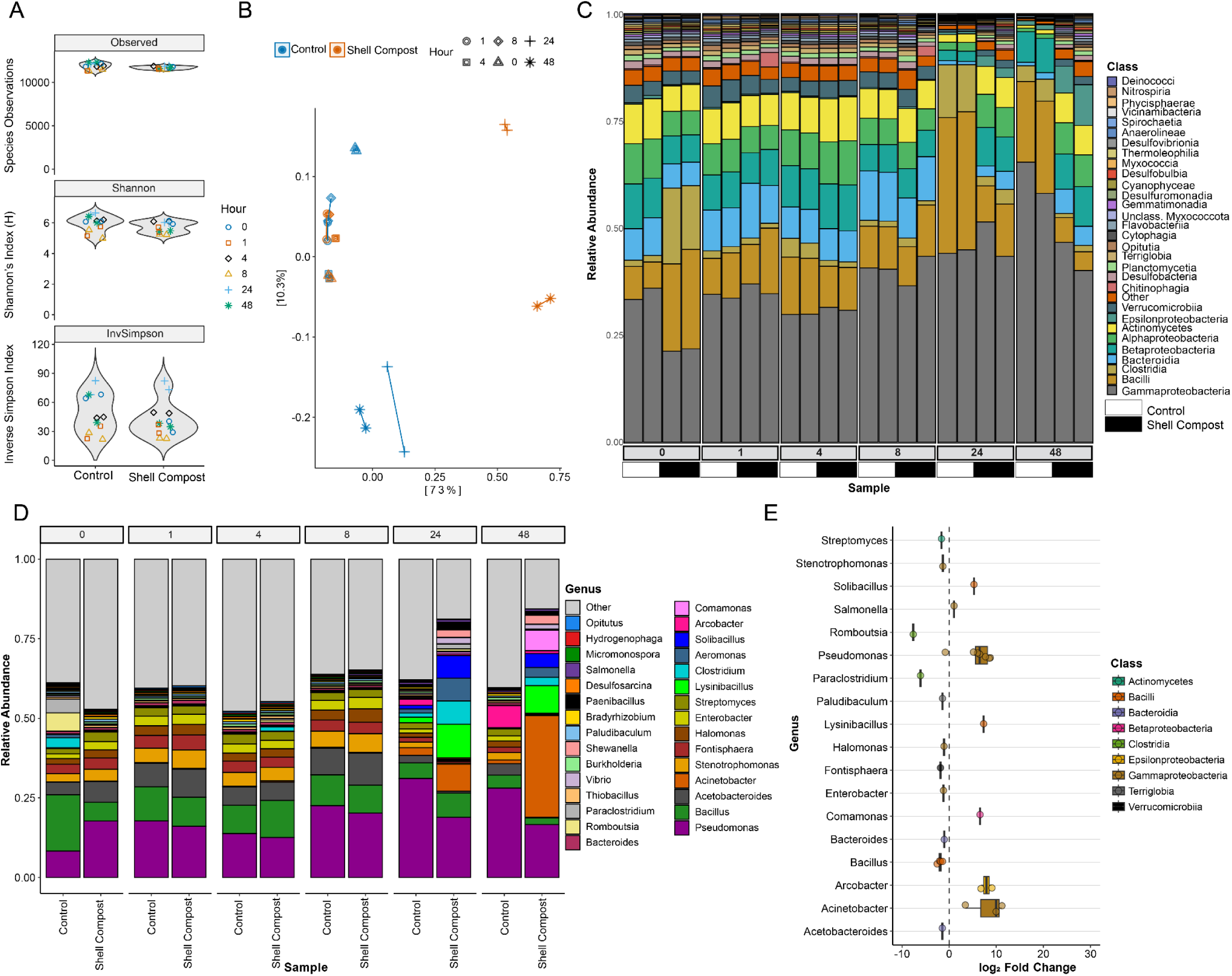
Metatranscript pool composition. (A) Violin plots showing alpha diversity between microbiome transcript profiling sets. (B) Principal coordinate analysis of microbial community transcript pool. (C) Stacked bar plot showing relative transcript read abundance of bacterial classes for each replicate. Black and white boxes are used to denote shell compost treated and control samples, respectively. (D) Stacked bar plot showing averaged relative transcript read abundance for bacterial genera. The top 30 most abundant genera are shown, while the rest are denoted as “Other” (colored gray). E) Boxplot showing likelihood ratios of genera among most abundant genera.

To identify microbial taxa actively growing in either treated or control soils, transcriptional abundance and taxonomic profiles of the *recA* gene were analyzed. This housekeeping gene was selected because it is constitutively expressed in metabolically active cells [56] and present in all microbial life. The protein domain encoding *recA* activity (PF00154) was not significantly enriched or depleted 48 hours after shell compost addition (p-value > 0.999), in line with the assumption that protein function is required for basic cellular maintenance and survival. Consistent with metagenomic and metatranscriptomic profiling, *recA* profiling indicated enrichment of the bacterial classes Gammaproteobacteria and Bacilli in the treated samples, while Betaproteobacteria, Alphaproteobacteria, Anaerolineae, and Espsilonproteobacteria were more abundant in the control samples (Fig. S2). It is concluded that the respective bacterial classes in each group were active members of the treated or control microbial communities.

### Significant Domain Enrichment 48 hours Post Shell Compost Addition

To identify which protein domains and associated functions, were selected and/or transcriptionally regulated in response to shell compost addition, phenotype-association analyses were performed (see Materials and Methods). Protein domains served as functional units to include all genes predicted to encode for a similar activity, regardless of taxonomy. As a result, it was possible to assess global functional changes throughout the microbiome. An increase in the relative abundance of a given protein domain at the DNA level indicates increased prevalence of organisms encoding this domain, suggesting selection for the corresponding function. An increase at the normalized RNA level indicates global transcriptional upregulation of genes harboring this domain across the community. These domains were designated as *functionally enriched* and corresponding transcripts considered *upregulated*. Conversely, domains more abundant in control samples were classified as *functionally depleted* in the treated condition and their corresponding transcripts designated *downregulated*.

Association analyses yielded 108 significantly functionally enriched domains with a p-value of 0 (Table S1). Notably, the glycoside hydrolase family 18 (GH18, PF00704) domain was significantly enriched, the very same domain in chitin degrading enzymes. Other functionally enriched domains included those involved in biosynthesis and metabolism. Transport, cell wall, and protein synthesis domains were also significantly enriched. Domains found in proteins involving stress response, resistance, and sporulation were also functionally enriched, suggesting community wide stress response post shell compost addition.

### Chitinase Gene Regulation

Domains found in enzymes related to chitin degradation were functionally enriched in samples collected 48 hours after shell compost addition. Transcripts containing these domains were encoded by a variety of bacterial taxa (Fig. 3A). Most transcripts were identified as containing a GH18 domain (EC 3.2.1.14, n=128), previously shown to have endochitinase, exochitinase, and N-acetylglucosaminidase activity [26]. Several upregulated transcripts containing GH18 were assigned to Gammaproteobacteria including *Aeromonas*, *Vibrio*, *Shewanella* spp., and members of the class Bacilli (Fig. 3A). Relative abundance of these taxa increased between initially sampled soils and soils treated with shell compost after 48 hours, contributing to reduced species diversity (Fig. 1). The GH19 domain, with known exochitinase activity [26], was less frequently detected (EC 3.2.1.14, n=19) and assigned to several bacteria, including both *Aeromonas* and *Vibrio* spp. (Fig. 3A). Several upregulated transcripts were also identified to encode GH20 domains (EC 3.2.1.52, n=43), annotated with beta-N-acetylglucosaminidase activity, catalyzing breakdown of diacetylchitobiose into N-acetylglucosamine [22].The abundance of transcripts from both identified and unidentified bacterial taxa indicates multiple microbial species contribute directly to chitin degradation (Fig. 3A).

**Figure 3:**
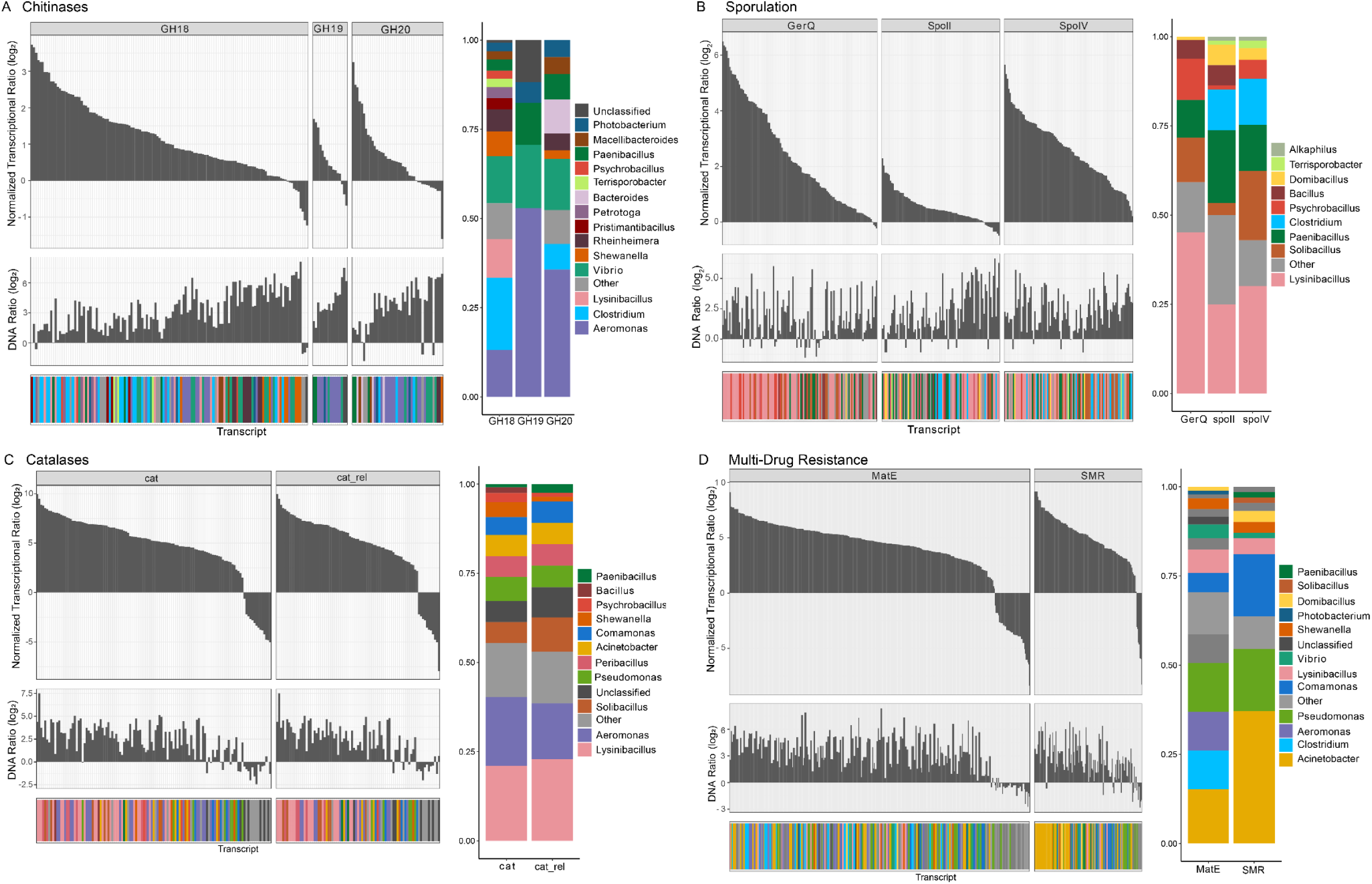
Upregulation community response to shell compost addition. Transcripts encoding select domains with significant enrichment 48 hours after chitin addition are depicted with normalized relative transcriptional changes, DNA ratios, and taxonomic assignment including those with (A) chitinase (B) sporulation (C) catalase and (D) multi-drug resistance activity. Normalized transcriptional changes are calculated by taking the ratio of relative RNA:DNA from shell compost treated to control samples at 48 hours, and DNA ratios = normalized DNA abundance in treatment/ normalized DNA abundance in control at 48 hours (see Methods).

### Regulation of Genes Related to Stress

Catalase function (EC 1.11.1.6) was significantly upregulated 48 hours after shell compost addition. Significant functional enrichment was observed for catalase (PF00199) and catalase-related immune-responsive domains (PF06628), both of which are involved in conversion of hydrogen peroxide to water and oxygen under oxidative stress. The most upregulated transcripts containing catalase domains were assigned to *Lysnibacillus* spp. and other Bacilli. However, several transcripts were also assigned to *Aeromonas*, *Pseudomonas*, *Acinetobacter*, and *Comamonas* spp. Despite being catalase positive, *Vibrio* spp. catalase transcripts were not detected (Fig. 3B).

Significant, functional enrichment of domains involved in diverse aspects of sporulation was also observed in samples collected 48 hours post shell compost addition. These domains included GerQ (PF09671), a spore coat protein produced by Bacillota, stage II sporulation protein SpoII (PF08486), involved in earlier steps of sporulation, and putative stage IV sporulation protein YqfD (PF06898), an essential protein for sporulation the exact function of which is unknown. Most upregulated transcripts containing these domains were assigned to classes Bacilli, and Clostridia (Fig. 3C). These community members are concluded to experience nutrient limitation and/or stress following shell compost amendment and subsequent changes in community structure.

Protein domains involved in multidrug resistance, Small Multidrug Resistance Protein (SMR, PF008930) and MatE (PF01554), were significantly enriched after shell compost addition. MatE-containing transcripts were taxonomically diverse, with over 50% of the upregulated transcripts assigned to *Acinetobacter*, *Clostridium*, *Aeromonas*, and *Pseudomonas*. Upregulated SMR transcripts derived mostly from *Acinetobacter*, *Pseudomonas*, and *Comamonas* spp. These taxa not only increased in relative abundance (Fig. 1) but also contributed substantially to the metatranscript pool post shell compost addition (Fig. 2).

### Significant Domain Depletion 48 hours after Shell Compost Addition

Similarly, 113 domains were significantly depleted in the shell compost treated samples at 48 hours. Several of these domains included various cytochrome systems, viral proteins, flagellar, and pili proteins. Significant depletion of domains central to various primary metabolism pathways was also observed. These domains included those with functional importance in carbon fixation, sulfur oxidation and reduction, nitrogen utilization, and carbohydrate metabolism (Table S2). Consistent with significant loss in species diversity 48 hours after chitin addition, depletion of domains found in various metabolic pathways highlights loss in functional diversity.

### Regulation of Genes Related Non-Carbon Based Metabolism

Transcripts containing domains involved in sulfur oxidation (Sox) were significantly downregulated in shell compost samples 48 hours after addition. SoxZ (PF08770) and SoxY (PF13501) domain containing proteins form critical complexes that are required for dissimilatory thiosulfate oxidation [57], an important process in sulfur cycling. Compared to the control, transcripts containing these domains were downregulated in treated samples, suggesting attenuation of this functional pathway, largely driven by a diverse, but poorly characterized group of taxa (Fig. 4A). Similar patterns were observed for transcripts containing AMO domains, that have ammonia and/or methane monooxygenase activity (Fig. 4D). Proteins containing AMO (PF02461) and AMOc (subunit C, PF04896) domains play key roles in nitrogen cycling and breakdown of chemically diverse hydrocarbons. Most of these transcripts derived from *Methylobacter* and *Methylomonas* spp., both of which are methanotrophs.

**Figure 4:**
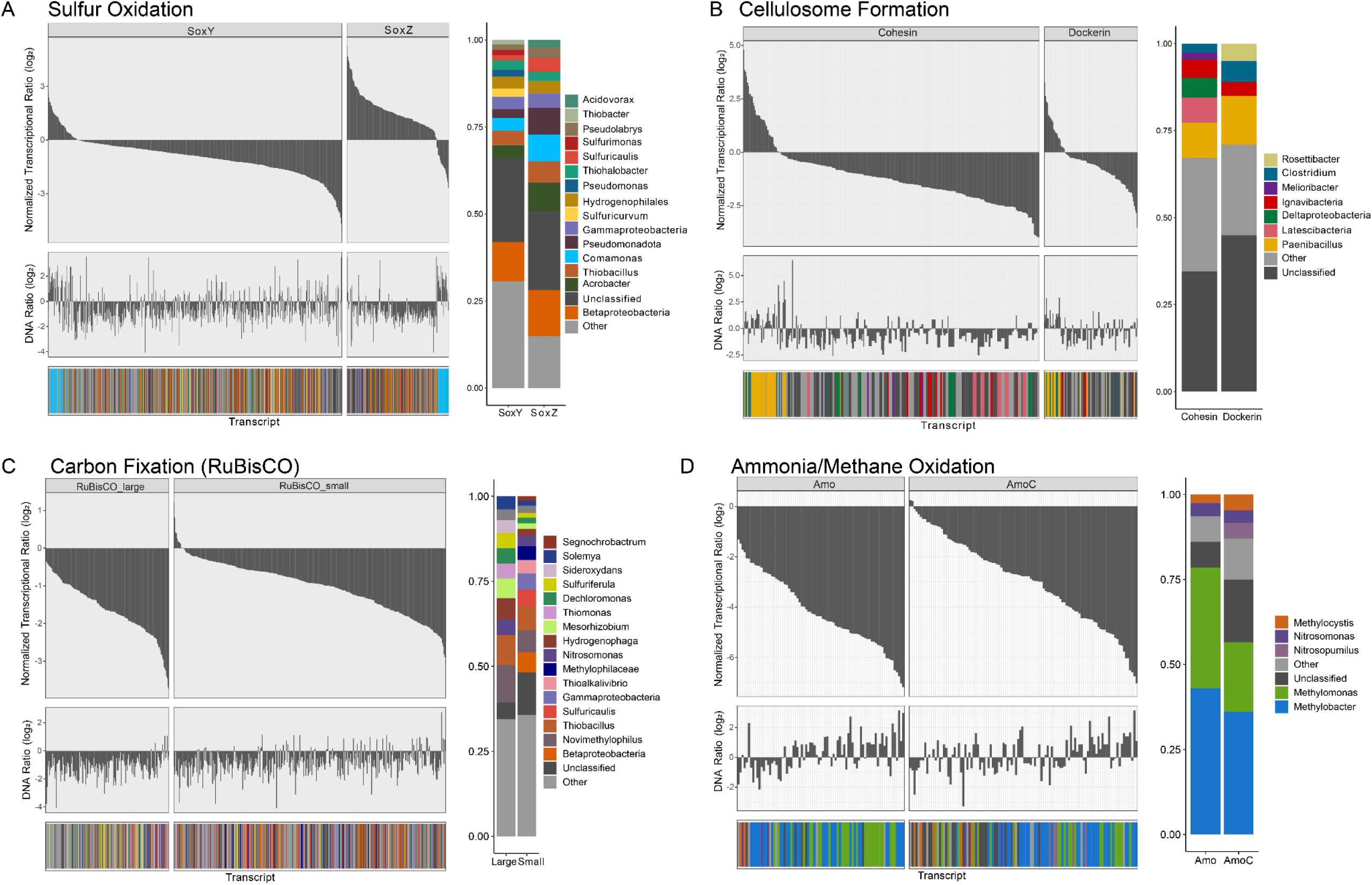
Downregulation community response to shell compost addition. Transcripts encoding select domains with significant depletion 48 hours after chitin addition are depicted with normalized relative transcriptional changes, DNA ratios, and taxonomic assignment including those with (A) sulfur oxidation (B) cellulosome formation (C) carbon fixation (RuBisCO) and (D) ammonia and methane oxidation activity. Normalized transcriptional changes are calculated by taking the ratio of relative RNA:DNA from shell compost treated to control samples at 48 hours, and DNA ratios = normalized DNA abundance in treatment/ normalized DNA abundance in control at 48 hours (see Methods).

### Regulation of Genes Related Carbon Fixation

Significant depletion of domains involved in other carbon metabolism pathways was also observed. Transcripts encoding predicted proteins with dockerin (PF00404) and cohesin (PF00963) domains were significantly downregulated 48 hours after shell compost addition. These proteins are required for an important interaction that enables formation of the cellulosome, a complex, multi-enzyme complex essential for cellulose degradation, the most abundant, naturally occurring polymer on Earth [58]. Similarly, transcripts encoding subunits of Ribulose-1,5-bisphosphate carboxylase/oxygenase (RuBisCO) were also downregulated. Both large (PF00016) and small (PF00101) subunits were depleted, indicating a functional loss in carbon fixation and photosynthetic processes.

### Detection of Genes related to Virulence and Antimicrobial Resistance

Given that several, potentially pathogenic taxa were enriched 24 hours after shell compost addition, a targeted analysis was used to detect presence and the expression of genes related to virulence and antimicrobial resistance. Several virulence factor (VF) genes were detected at 24 and 48 hours (Fig. 5, S2). Their functions varied, but included those involved in adherence, effector delivery, and motility. Most VFs detected were from *Aeromonas* and *Vibrio* spp. (Fig. 5A,B, S2). Several genes encoding exotoxins were also detected including *aerA* encoding aerolysin A and *rtx* genes that encode rtx toxins (*Aeromonas*), *hlyA* encoding hemolysin (*Vibrio* spp.), and several *Bacillus* exotoxin genes including *alo* which encodes anthrolysin. A smaller subset of VFs was found to be actively expressed 24 and 48 hours after shell compost addition (Fig. 6A, 6B, S2). Several of the most highly expressed transcripts were involved in effector delivery from *Acinetobacter*, *Aeromonas*, and *Vibrio* spp.. Only one exotoxin gene, *aerA*, showed detectable transcriptional levels (Fig. 6).

**Figure 5:**
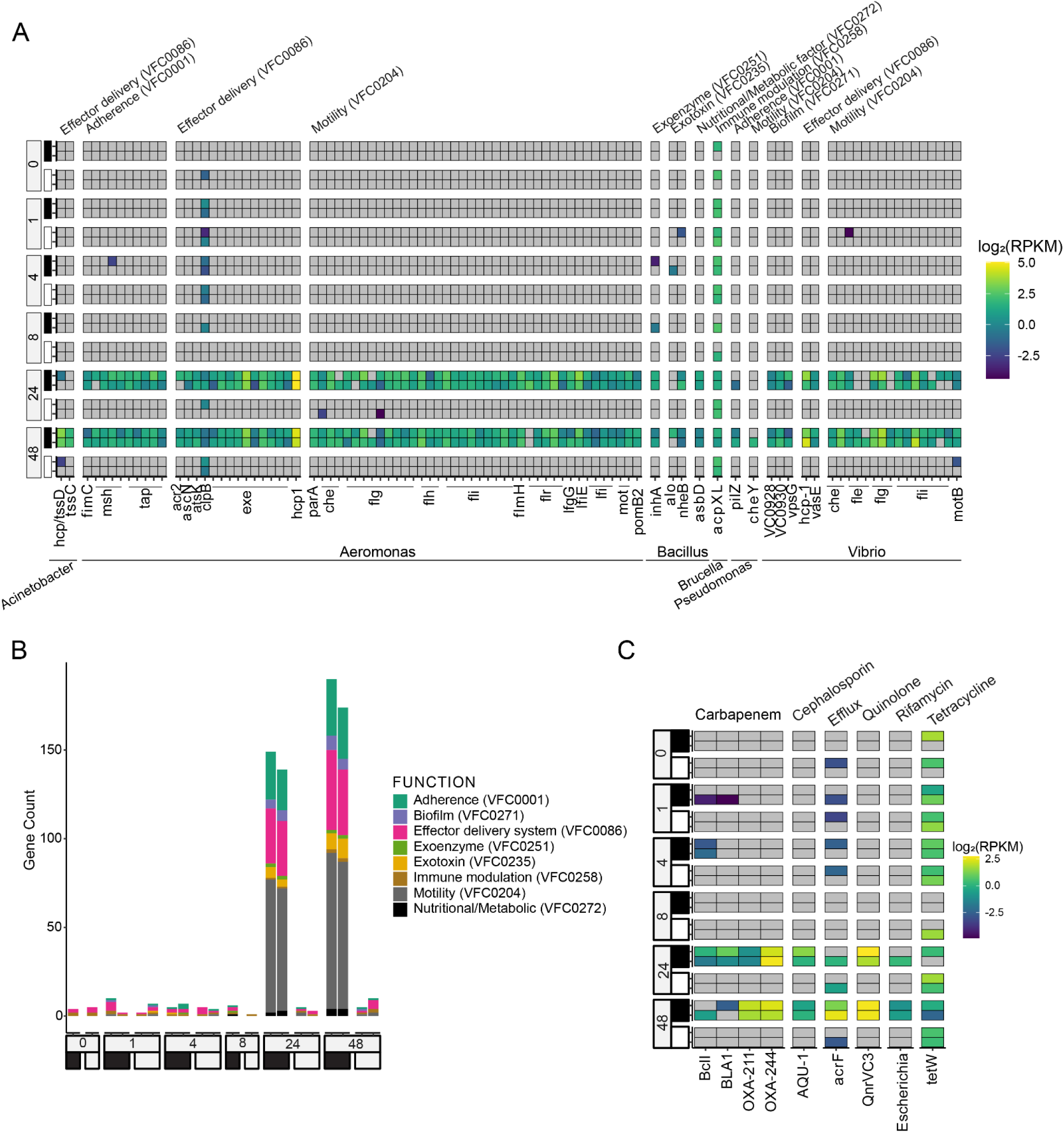
Microbiome profiling for virulence factor determinants and antimicrobial resistance gene abundance. (A) Detection and relative abundance of virulence factor genes where RPKM ≥2 in at least one sample. (B) Stacked bar plots showing gene count for each replicate are displayed. (C) Relative gene abundance of detected antimicrobial resistance genes. Black and white boxes are used to denote shell compost treated and control samples, respectively.

**Figure 6:**
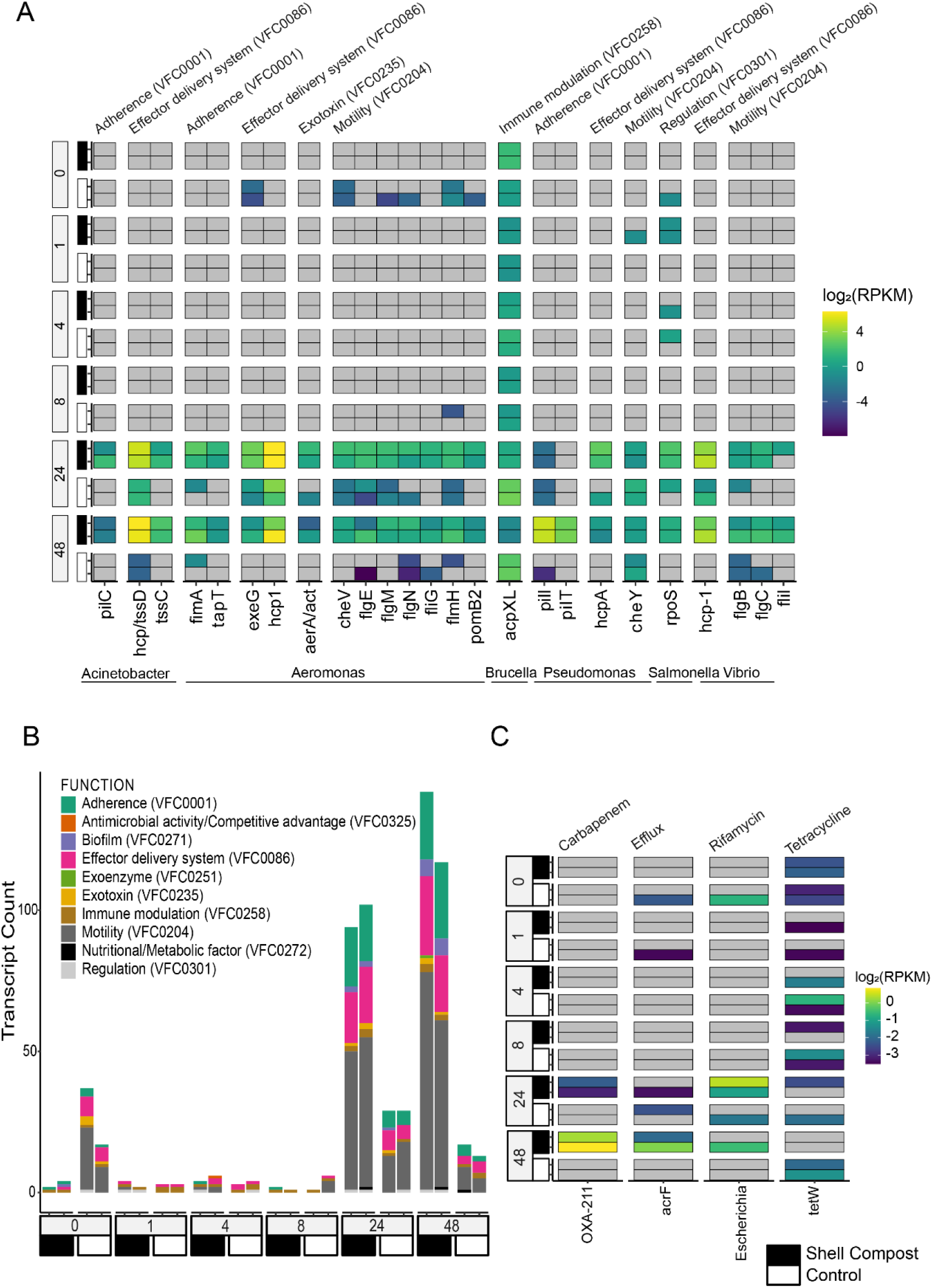
Microbiome profiling for virulence factor determinants and antimicrobial resistance gene expression. (A) Detection and relative expression of virulence factor genes where RPKM ≥2 in at least one sample. (B) Stacked bar plots showing relative transcript count for each replicate are displayed. (C) Relative gene expression of detected antimicrobial resistance genes Black and white boxes are used to denote shell compost treated and control samples, respectively.

Presence and expression of genes involved in antimicrobial resistance (ARGs) were also assessed. Relative abundance of these genes was lower than VF genes (Fig. 5C). Tetracycline resistance gene *tetW* was detected in nearly every sample, regardless of treatment, exhibiting low, but detectable, levels of expression (Fig. 5C, 6C). At 24 and 48 hours post shell compost addition, several genes responsible for carbapenem resistance were detected (Fig. 5C). Expression of ARGs was sporadic, with OXA-211 showing greatest relative expression 48 hours after shell compost addition (Fig. 6C). While several ARGs were detected, they were upregulated to a lesser extent compared to VFs (Fig. 5C, 6C).

## Discussion

As native environments continue to be altered by nutrient loading, the microbial community dynamics of these systems need to be investigated to understand ecosystem function and resilience. Employing a natural, chitin-rich substrate (shell compost), allowed demonstration of the effects of a single pulse event in selectively enriching for chitin degradation and selection for pathogenic taxa naturally existing in coastal ecosystems. Dynamic shifts in transcript abundance revealed not only an increase in chitin degradation, as was expected, but also stress and virulence responses (Figs. 3,5,6). While metagenomic analysis revealed a marked increase in *Aeromonas*, *Vibrio*, and *Shewanella* spp., other bacterial genera showing modest increase in abundance, e.g. *Acinetobacter* and *Clostridium* spp., were some of the most transcriptionally active community members within the amended soils (Fig. 1,2). These findings underscore how nutrient pulses can rapidly alter microbial communities by enriching for selected species, some of which are potential pathogens [11, 18, 59].

In this study it was hypothesized that shell compost addition provided a nutrient rich substrate that would stimulate migration and biofilm formation. Ultimately, this resulted in functional shift toward virulence and resistance, potentially to combat competition (Fig. 7). While the hypothesized upregulation of chitin degrading genes was observed, significant upregulation in genes related to stress response and resistance were also detected. Upregulation of catalase genes across many bacterial taxa suggests a community wide response to oxidative stress. Similarly, sporulation genes were upregulated, indicating nutrient limitation and/or stress in the Bacillota bacteria despite nutrient replete conditions. These bacteria appear to have adopted this regulated survival mechanism, considered a last resort under unfavorable environmental conditions [60–62], as an alternative survival strategy under highly competitive settings.

**Figure 7:**
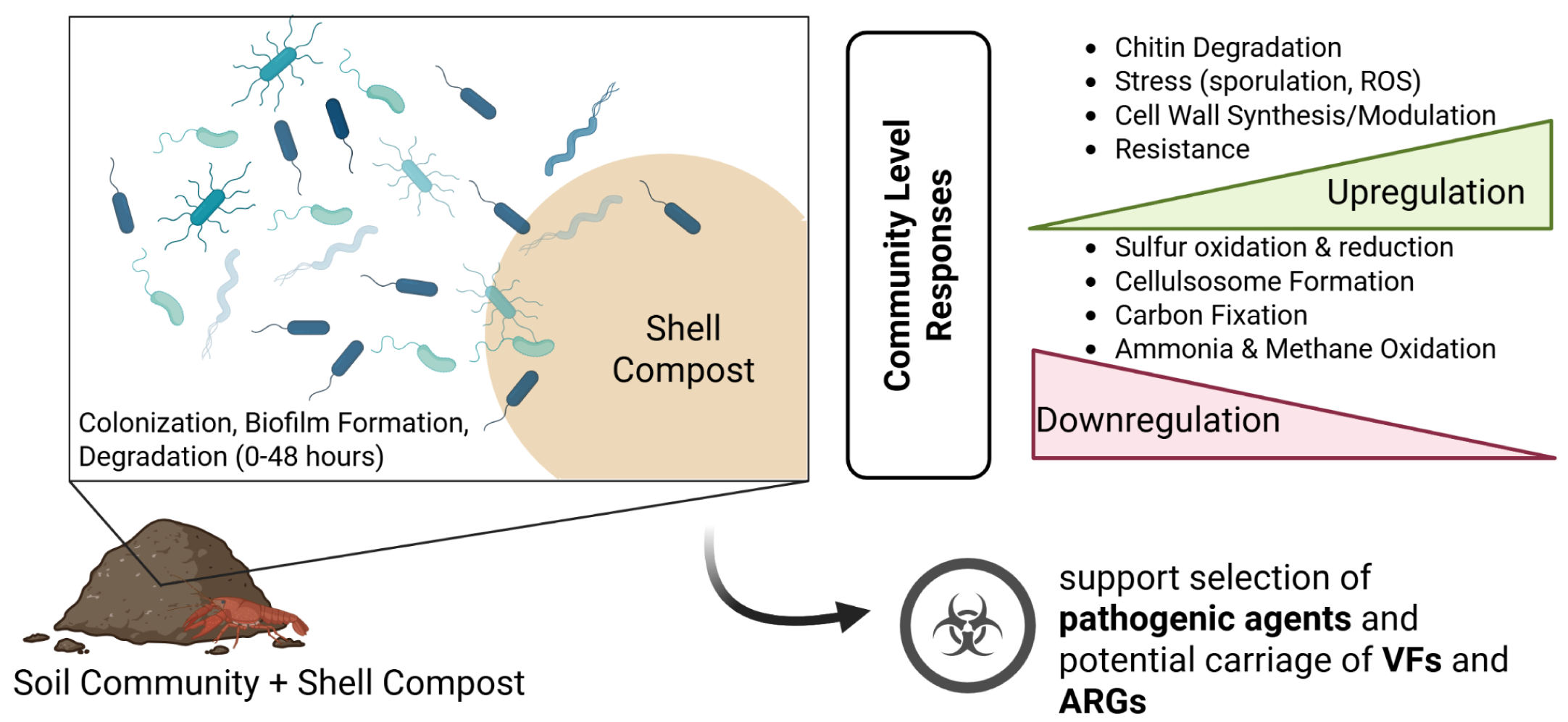
Schematic overview of microbiome response to shell compost addition. Community succession takes place in as little as 48 hours, resulting in the upregulation and downregulation of several functionally important enzymes. At 48 hours, the detection of potentially pathogenic taxa increases, in addition to the detection and relative expression of virulence factors (VFs) and antibiotic resistance genes (ARGs), with potential implications for human health.

Similarly, significant upregulation of genes involved in multi-drug resistance suggests active competition among community members for the chitin-rich substrate. Targeted analysis for a broad panel of VFs supports this notion as genes for motility, biofilm formation/adherence, and effector delivery were not only enriched 24-48 hours after chitin addition, but were also detectably expressed as well. *Aeromonas* and *Vibrio* spp., were responsible for a large portion of the detected and expressed VF genes. As a result, biodiversity was reduced within the microbial community, resulting in loss of functionally important traits that drive biogeochemical cycling, evidenced by downregulation of genes involved in carbon fixation, sulfur, and nitrogen metabolism (Fig. 1, 4, 5, 6, Table S2).

These results contribute to a growing body of work highlighting the enrichment of microbial species, some of which are potential pathogens, in northern latitudes under changing climate and nutrient loading scenarios. These findings indicate some of these enriched taxa are also resistant to antibiotics, a growing concern in natural systems worldwide [63]. Extensive work has shown increased temperature, change in salinity and pH, and extreme events such as flooding, can stimulate proliferation of microbial species, including potential pathogens, in coastal ecosystems [13, 17]. The importance of chitin-rich substrates in aquatic ecosystems includes providing a critical source of nutrients and promotion of biofilm formation for taxa, some of which may be human pathogens. Other research has demonstrated nutrient influx and the presence of shellfish in a coastal environment provide substrates suitable for biofilm formation, a function related to virulence [64–67]. Thus, the distribution of chitin-rich substrates can have important predictive and public health implications for coastal communities.

The results of this study highlight the lack of resilience of coastal communities where within 48 hours, a significant loss of microbial species diversity and concurrent downregulation of metabolic genes that several biogeochemical cycles can be observed (Table 2, Fig. 5,6). This highlights the potential for certain bacterial taxa, which naturally occur in low abundance, to rapidly dominate the microbial community in response to chitin-rich substrates. The findings presented here are noteworthy given that samples were collected mid-October in northern Massachusetts, at a time and location not often associated with vibriosis and other pathogen outbreak [11, 68]. As changes in weather patterns and extreme events intensify, in addition to nutrient loading, the results of this study underscore the importance of microbial surveillance of coastal systems which may harbor potentially pathogenic bacteria that can significantly alter a microbiome’s ability contribute to biogeochemical cycling and associated important ecosystem services.

## Data Availability

Raw reads are available on the NCBI Sequencing Read Archive (SRA) under BioProject: PRJNA1293438.

## Acknowledgements

We thank the NEB Sequencing Core and NEB IT Department for their assistance in sequencing and technical support. We are also grateful to Andy Ge for computational and bioinformatic support.

