## Supplementary material for "Microbial Community multi-omic analysis of marsh sediment post crustacean shell compost enrichment: pathogen emergence and community response": Table S; Fig. S

### Supplemental Figures and Tables

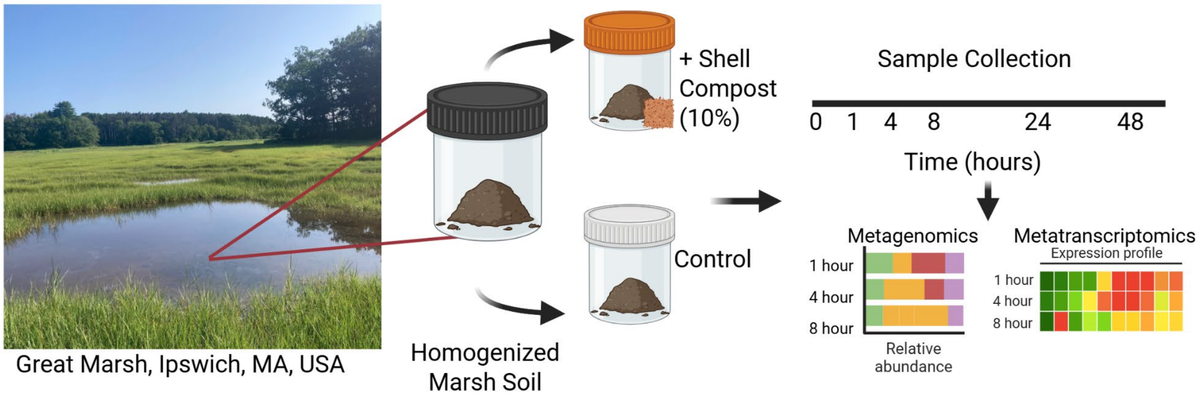


**Figure S1:** Schematic overview of the microcosm experiments and sample collection. Figure generation was completed with Biorender.

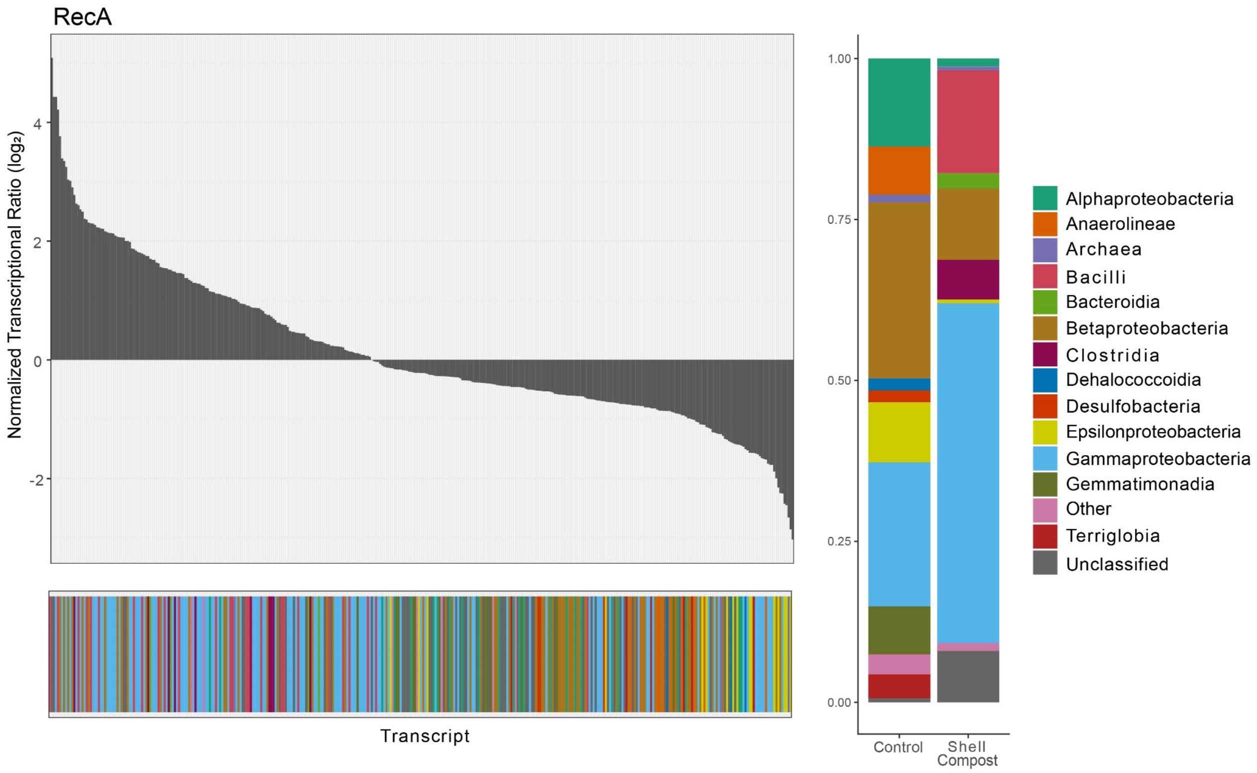


**Figure S2:** Metatranscriptomic profiling of recA gene A) Normalized transcriptional changes of housekeeping gene recA at 48 hours after chitin addition. B) Relative transcript abundance of bacterial classes in the control and treated microcosms.

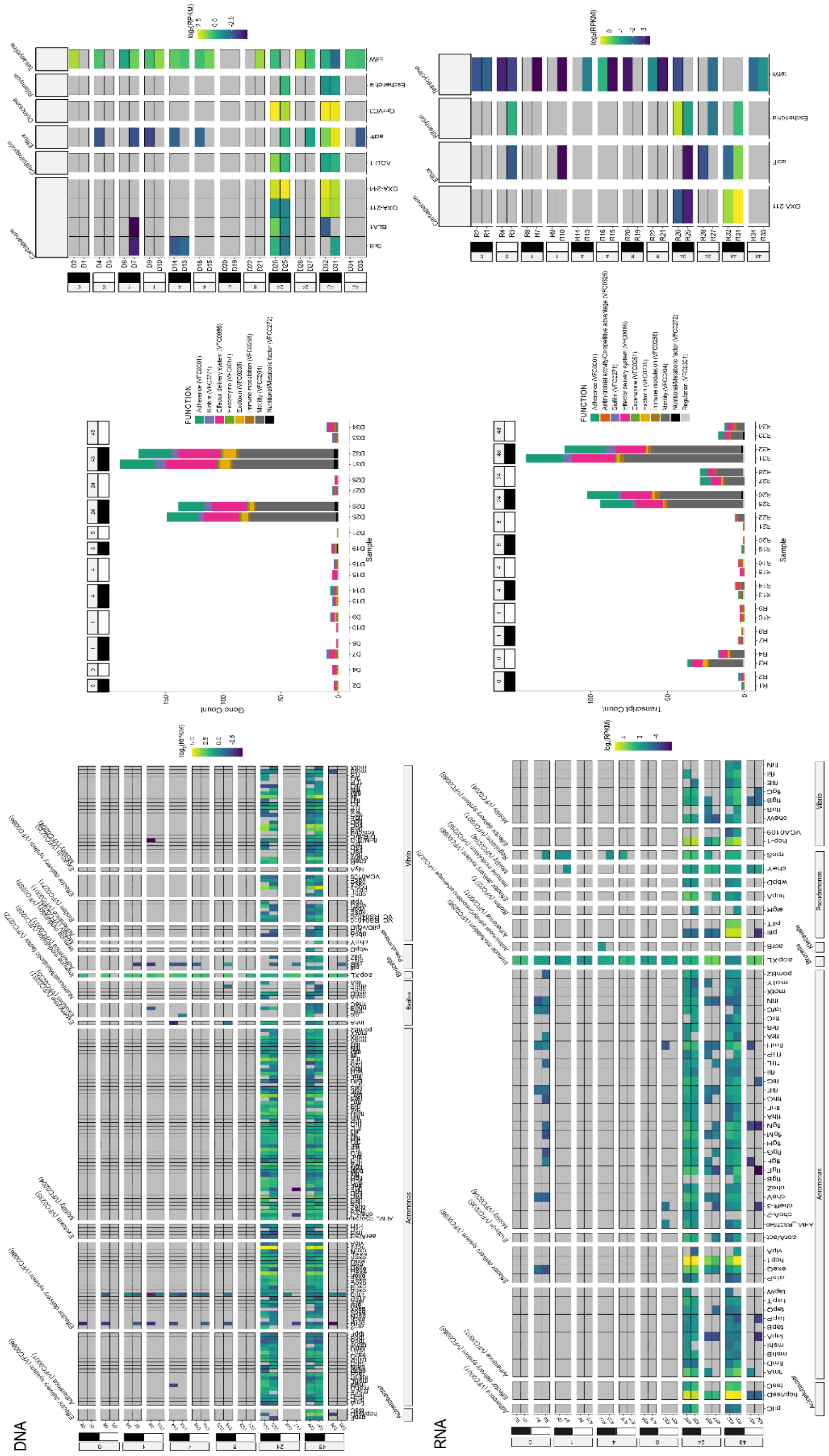


**Figure S3:** Microbiome profiling for virulence factor determinants and antimicrobial resistance. A) Detection and relative abundance of virulence factor genes where RPKM ≥1 in at least one sample. Stacked bar plots showing gene count for each replicate are also displayed. B) Detection and relative expression of virulence factor genes. Stacked bar plots showing transcript count for each replicate are also displayed. C) Relative gene abundance and expression of detected antimicrobial resistance genes

**Table S1:** List of significantly enriched protein domains 48 hours after chitin addition (p-value <0.001).

| PFAM | Name | Description | Transcript Count in Treated | Total Transcripts |
|---|---|---|---|---|
| PF00501.31 | AMP-binding | AMP-binding | 1223 | 1831 |
| PF02875.24 | Mur_ligase_C | Muramyl ligase, glutamate ligase | 238 | 291 |
| PF01039.25 | Carboxyl_trans | Carboxyl transferase | 388 | 512 |
| PF00849.25 | PseudoU_synth_2 | RNA pseudouridylate synthase | 352 | 465 |
| PF06628.15 | Catalase-rel | Catalase related immune responsive | 118 | 134 |
| PF00165.26 | HTH_AraC | Bacterial regulatory helix-turn-helix, AraC | 328 | 432 |
| PF00155.24 | Aminotran_1_2 | Aminotransferase class I and II | 970 | 1346 |
| PF16113.8 | ECH_2 | Enoyl-CoA hydratase/isomerase | 799 | 1080 |
| PF04069.15 | OpuAC | Substrate binding domain of ABC-type glycine betaine transport system | 142 | 155 |
| PF13302.10 | Acetyltransf_3 | Acetyltransferase (GNAT) | 173 | 214 |
| PF04851.18 | ResIII | Type III restriction enzyme, res subunit | 207 | 253 |
| PF03401.17 | TctC | Tripartite tricarboxylate transporter family receptor | 610 | 760 |
| PF02028.20 | BCCT | Betaine, carnitine, choline family transporter | 107 | 115 |

| | | | | |
|---|---|---|---|---|
| PF02824.24 | TGS | TGS domain | 220 | 272 |
| PF03180.17 | Lipoprotein_9 | NlpA lipoprotein | 101 | 110 |
| PF00198.26 | 2-oxoacid_dh | 2-oxoacid dehydrogenase (catalytic) | 455 | 634 |
| PF04095.19 | NAPRTase | Nicotinate phosphorribosyltransferase (NAPRTase) family | 65 | 67 |
| PF00199.22 | Catalase | catalase | 182 | 202 |
| PF00271.34 | Helicase_C | Helicase conserved C-terminal | 631 | 831 |
| PF20579.1 | LapA | LapA | 110 | 119 |
| PF02463.22 | SMC_N | RecF/RecN/SMC N terminal domain | 344 | 443 |
| PF01266.27 | DAO | FAD dependent oxidoreductase | 654 | 933 |
| PF01554.21 | MatE | multidrug/oligosaccharidyl-lipid/polysaccharide (MOP) flippase superfamily | 138 | 159 |
| PF00133.25 | tRNA-synt_1 | tRNA synthetases class I (I, L, M and V) | 498 | 691 |
| PF01225.28 | Mur_ligase | muramyl ligase | 182 | 229 |
| PF08028.14 | Acyl-CoA_dh_2 | Acyl-CoA dehydrogenase, C-terminal | 528 | 710 |
| PF00275.23 | EPSP_synthase | EPSP synthase (3-phosphoshikimate 1-carboxyvinyltransferase) | 230 | 302 |
| PF03553.17 | Na_H_antiporter | Na+/H+ antiporter family | 134 | 143 |

| | | | | |
|---|---|---|---|---|
| PF02786.20 | CPSase_L_D2 | Carbamoyl-phosphate synthase L chain, ATP binding | 559 | 734 |
| PF02378.21 | PTS_EIIC | Phosphotransferase system, EIIC | 191 | 192 |
| PF00324.24 | AA_permease | amino acid permease | 381 | 452 |
| PF00083.27 | Sugar_tr | sugar and other transport | 314 | 386 |
| PF02222.25 | ATP-grasp | ATP grasp | 243 | 310 |
| PF09680.13 | YjcZ_2 | family of unknown function | 68 | 69 |
| PF09334.14 | tRNA-synt_1g | tRNA synthetases class I (M) | 426 | 566 |
| PF01432.23 | Peptidase_M3 | Peptidase family M3 | 203 | 251 |
| PF00015.24 | MCPsignal | Methyl-accepting chemotaxis protein (MCP) signaling | 899 | 1230 |
| PF00725.25 | 3HCDH | 3-hydroxyacyl-CoA dehydrogenase, C-terminal | 372 | 487 |
| PF02771.19 | Acyl-CoA_dh_N | Acyl-CoA dehydrogenase, N-terminal | 840 | 1166 |
| PF09917.12 | DUF2147 | Domain of unknown function | 65 | 67 |
| PF00528.25 | BPD_transp_1 | Binding-protein-dependent transport system inner membrane component | 1029 | 1482 |
| PF00367.23 | PTS_EIIB | phosphotransferase system, EIIB | 87 | 88 |
| PF00905.25 | Transpeptidase | Penicillin binding protein transpeptidase | 273 | 362 |

| | | | | |
|---|---|---|---|---|
| PF03741.19 | TerC | heavy metal resistance | 166 | 189 |
| PF00126.30 | HTH_1 | Bacterial regulatory helix-turn-helix protein, lysR | 984 | 1217 |
| PF12697.10 | Abhydrolase_6 | Alpha/beta hydrolase family | 403 | 536 |
| PF00395.23 | SLH | S-layer homology protein | 608 | 652 |
| PF00171.25 | Aldedh | Aldehyde dehydrogenase family | 1511 | 2061 |
| PF00893.22 | Multi_Drug_Res | Small Multidrug Resistance protein | 84 | 91 |
| PF20439.1 | SpoIVA_C | Sporulation stage IV protein A, C-terminal | 79 | 80 |
| PF02342.21 | TerD | stress response in multiple metal recognition complexes | 74 | 78 |
| PF02421.21 | FeoB_N | Ferrous iron transport protein B | 281 | 365 |
| PF00860.23 | Xan_ur_permease | permease family | 269 | 286 |
| PF08264.16 | Anticodon_1 | Anticodon-binding domain of tRNA ligase | 306 | 380 |
| PF12833.10 | HTH_18 | Helix-turn-helix | 480 | 643 |
| PF00441.27 | Acyl-CoA_dh_1 | Acyl-CoA dehydrogenase, C-terminal | 992 | 1321 |
| PF13336.9 | AcetylCoA_hyd_C | Acetyl-CoA hydrolase, C-terminal | 162 | 194 |
| PF00664.26 | ABC_membrane | ABC transporter transmembrane region | 232 | 298 |
| PF00106.28 | adh_short | short chain dehydrogenase | 1302 | 2091 |

| | | | | |
|---|---|---|---|---|
| PF00440.26 | TetR_N | tetracycline resistance repressor | 496 | 626 |
| PF20438.1 | SpoIVA_middle | Stage IV sporulation protein A, middle | 95 | 96 |
| PF17876.4 | CSD2 | cold shock | 96 | 107 |
| PF00465.22 | Fe-ADH | Iron-containing alcohol dehydrogenase | 343 | 431 |
| PF01810.21 | LysE | LysE type translocator | 264 | 310 |
| PF08245.15 | Mur_ligase_M | muramyl ligase middle | 339 | 421 |
| PF00561.23 | Abhydrolase_1 | Alpha/beta hydrolase family | 681 | 982 |
| PF00724.23 | Oxidored_FMN | NADH:flavin oxidoreductase | 247 | 303 |
| PF00289.25 | Biotin_carb_N | Biotin carboxylase, N-terminal | 300 | 399 |
| PF09547.13 | SpoIVA_ATPase | Stage IV sporulation protein A, ATPase | 105 | 106 |
| PF05175.17 | MTS | Methyltransferase small domain | 204 | 248 |
| PF01926.26 | MMR_HSR1 | 50S ribosome-binding GTPase | 804 | 1164 |
| PF03922.17 | OmpW | outer membrane protein W | 115 | 132 |
| PF02817.20 | E3_binding | e3 binding domain | 389 | 527 |
| PF02785.22 | Biotin_carb_C | Biotin carboxylase C-terminal | 297 | 374 |
| PF00392.24 | GntR | Bacterial regulatory proteins, gntR family | 422 | 550 |

| | | | | |
|---|---|---|---|---|
| PF12806.10 | Acyl-CoA_dh_C | Acetyl-CoA dehydrogenase, C-terminal like | 208 | 247 |
| PF12146.11 | Hydrolase_4 | Serine aminopeptidase, S33 | 541 | 724 |
| PF00990.24 | GGDEF | Diguanylate cyclase, GGDEF domain | 835 | 1261 |
| PF00364.25 | Biotin_lipoyl | Biotin-requiring enzyme | 748 | 1062 |
| PF00206.23 | Lyase_1 | fumarate lyase | 374 | 510 |
| PF00108.26 | Thiolase_N | Thiolase, N-terminal | 738 | 1123 |
| PF13685.9 | Fe-ADH_2 | Iron-containing alcohol dehydrogenase | 291 | 361 |
| PF00892.23 | EamA | EamA-like transporter family | 340 | 449 |
| PF02737.21 | 3HCDH_N | 3-hydroxyacyl-CoA dehydrogenase NAD binding | 384 | 514 |
| PF13603.9 | tRNA-synt_1_2 | Leucyl-tRNA synthetase | 201 | 252 |
| PF00270.32 | DEAD | DEAD/DEAH box helicase | 509 | 692 |
| PF13193.9 | AMP-binding_C | AMP-binding enzyme, C-terminal domain | 727 | 1108 |
| PF13520.9 | AA_permease_2 | Amino acid permease | 386 | 464 |
| PF02770.22 | Acyl-CoA_dh_M | Acyl-CoA dehydrogenase, middle domain | 896 | 1202 |
| PF03466.23 | LysR_substrate | LysR substrate binding domain | 1059 | 1311 |
| PF00587.28 | tRNA-synt_2b | tRNA synthetase class II core domain | 377 | 531 |

| | | | | |
|---|---|---|---|---|
| PF10612.12 | Spore-coat_CotZ | Spore coat protein Z | 62 | 63 |
| PF01546.31 | Peptidase_M20 | Peptidase family M20/M25/M40 | 403 | 550 |
| PF00375.21 | SDF | Sodium:dicarboxylate symporter family | 163 | 176 |
| PF13245.9 | AAA_19 | AAA+ family 19 | 162 | 200 |
| PF00672.28 | HAMP | HAMP domain | 468 | 585 |
| PF00005.30 | ABC_tran | ABC transporter | 2821 | 4342 |
| PF02729.24 | OTCace_N | Aspartate/ornithine carbamoyltransferase | 249 | 329 |
| PF00378.23 | ECH_1 | Enoyl-CoA hydratase/isomerase | 853 | 1175 |
| PF03061.25 | 4HBT | Thioesterase superfamily | 322 | 429 |
| PF17936.4 | Big_6 | Bacterial Ig domain | 112 | 117 |
| PF01391.21 | Collagen | Collagen triple helix repeat (20 copies) | 209 | 245 |
| PF00704.31 | Glyco_hydro_18 | glycosyl hydrolase family 18 | 120 | 124 |
| PF13304.9 | AAA_21 | AAA domain, family 21 | 623 | 868 |
| PF00269.23 | SASP | Small, acid-soluble spore proteins, | 88 | 89 |
| PF00202.24 | Aminotran_3 | Aminotransferase class-III | 685 | 970 |
| PF07690.19 | MFS_1 | Major Facilitator Superfamily | 1026 | 1447 |

| | | | | |
|---|---|---|---|---|
| PF06434.16 | Aconitase_2_N | Aconitate hydratase 2 N-terminus | 175 | 213 |

**Table S2:** List of significantly depleted protein domains 48 hours after chitin addition (p-value <0.001).

| PFAM | Name | Description | Transcript Count in Control | Total Transcripts |
|---|---|---|---|---|
| PF03646.18 | FlaG | flagellar protein | 97 | 233 |
| PF00016.23 | RuBisCO_large | Rubisco large | 142 | 252 |
| PF00978.24 | RdRP_2 | RNA dependent RNA polymerase | 50 | 74 |
| PF00404.21 | Dockerin_1 | Dockerin in cellulosome | 129 | 234 |
| PF02085.19 | Cytochrom_CIII | Class III cytochrome C family | 163 | 542 |
| PF09699.13 | Paired_CXXCH_1 | Doubled CXXCH motif | 396 | 1075 |
| PF04969.19 | CS | heat shock recruitment | 108 | 222 |
| PF09242.14 | FCSD-flav_bind | Flavocytochrome C sulfide dehydrogenase, flavin binding | 46 | 82 |
| PF17886.4 | ArsA_HSP20 | heat shock like protein found in ArsA | 679 | 1807 |
| PF02123.19 | RdRP_4 | Viral RNA-directed RNA polymerase | 49 | 87 |
| PF18962.3 | Por_Secre_tail | Secretion system C-terminal sorting domain | 523 | 885 |
| PF00428.22 | Ribosomal_60s | 60s Acidic ribosomal protein | 25 | 31 |
| PF13372.9 | Alginate_exp | alginate export | 76 | 128 |
| PF02472.19 | ExbD | Biopolymer transport protein | 186 | 679 |

| | | | | |
|---|---|---|---|---|
| PF08770.14 | SoxZ | Sulfur oxidation protein | 118 | 232 |
| PF04964.17 | Flp_Fap | fimbriae associated protein in pilin | 327 | 437 |
| PF13715.9 | CarbopepD_reg_2 | carboxypepD_reg-like domain | 551 | 1379 |
| PF00593.27 | TonB_dep_Rec | TonB dependent outer membrane receptor | 451 | 1580 |
| PF18046.4 | FKBP26_C | protein folding | 48 | 83 |
| PF14522.9 | Cytochrome_C7 | Cytochrome C7 and related cytochrome C | 491 | 1458 |
| PF08206.14 | OB_RNB | Ribonuclease B OB domain | 321 | 918 |
| PF02201.21 | SWIB | putative p53 associated protein | 55 | 107 |
| PF16371.8 | MetallophosN | Calcineurin-like phosphoesterase, N-terminal | 40 | 53 |
| PF13937.9 | DUF4212 | Domain of unknown function | 83 | 189 |
| PF07596.14 | DUF1559 | Domain of unknown function | 144 | 207 |
| PF02433.18 | FixO | Cytochrome C oxidase, mono-heme subunit | 146 | 348 |
| PF00114.22 | Pilin | pilin bacterial filament | 236 | 414 |
| PF00313.25 | CSD | Cold shock DNA-binding protein | 530 | 1271 |
| PF09430.13 | EMC7_beta-sandw | ER membrane protein complex subunit 7, beta-sandwich | 78 | 172 |
| PF12139.11 | APS-reductase_C | Adenosine-5'-phosphosulfate reductase beta subunit | 170 | 382 |

| | | | | |
|---|---|---|---|---|
| PF00120.27 | Gln-synt_C | glutamate synthetase, catalytic domain | 256 | 913 |
| PF00808.26 | CBFD_NFYB_HMF | Histone-like transcription factor (CBF/NF-Y) and archaeal histone | 32 | 36 |
| PF07980.14 | SusD_RagB | starch utilization system for glycan uptake | 97 | 263 |
| PF05538.14 | Campylo_MOMP | Campylobacter major outermembrane protein | 65 | 88 |
| PF07963.15 | N_methyl | 2-oxoadipate dioxygenase/decarboxylase | 855 | 1736 |
| PF13609.9 | Porin_4 | Gram negative porin | 958 | 1576 |
| PF00011.24 | HSP20 | heat shock 20 family | 748 | 2039 |
| PF01077.25 | NIR_SIR | Nitrite and sulphite reductase 4Fe-4S | 254 | 649 |
| PF00101.23 | RuBisCO_small | Rubisco small | 69 | 146 |
| PF13620.9 | CarboxypepD_reg | Carboxypeptidase regulatory-like | 630 | 1907 |
| PF09361.13 | Phasin_2 | phasin protein, granule associated | 157 | 419 |
| PF18201.4 | PIH1_CS | PIH1 CS-like domain | 43 | 53 |
| PF07715.18 | Plug | TonB-dependent receptor plug domain | 540 | 1721 |
| PF13570.9 | PQQ_3 | Beta-alanine-activating enzyme, beta-propeller | 183 | 593 |
| PF11839.11 | Alanine_zipper | Alanine-zipper, major outer membrane lipoprotein | 63 | 136 |
| PF13442.9 | Cytochrome_CBB3 | Cytochrome C oxidase, cbb3-type, subunit III | 581 | 2323 |

| | | | | |
|---|---|---|---|---|
| PF14686.9 | fn3_3 | Polysaccharide lyase family 4, domain II | 128 | 271 |
| PF00963.21 | Cohesin | Cohesin interaction for cellulosome formation | 117 | 196 |
| PF00022.22 | Actin | Actin | 129 | 377 |
| PF04077.15 | DsrH | DsrH like protein | 68 | 165 |
| PF09694.13 | Gcw_chp | Conserved bacterial protein of unknown function | 177 | 263 |
| PF18205.4 | VPDSG-CTERM | VPDSG-CTERM motif | 87 | 100 |
| PF20071.2 | DUF6467 | Domain of unknown function | 1141 | 2786 |
| PF00729.21 | Viral_coat | viral coat protein S domain | 75 | 113 |
| PF00496.25 | SBP_bac_5 | Bacterial extracellular solute-binding protein | 541 | 2125 |
| PF00543.25 | P-II | Nitrogen regulatory protein P-II | 344 | 749 |
| PF02788.19 | RuBisCO_large_N | Rubisco large N terminal | 108 | 179 |
| PF04744.15 | Monooxygenase_B | Monooxygenase subunit B protein | 73 | 77 |
| PF00680.23 | RdRP_1 | Viral RNA-dependent RNA polymerase | 140 | 259 |
| PF14881.9 | Tubulin_3 | Tubulin | 25 | 27 |
| PF13860.9 | FlgD_ig | FlgD Ig-like domain | 379 | 833 |
| PF01011.24 | PQQ | Glucose/ethanol/alcohol dehydrogenase, beta-propeller domain | 223 | 751 |

| | | | | |
|---|---|---|---|---|
| PF13458.9 | Peripla_BP_6 | periplasmic binding protein | 500 | 2211 |
| PF00125.27 | Histone | Core histone H2A/H2B/H3/H4 domain | 47 | 52 |
| PF03460.20 | NIR_SIR_ferr | Nitrite/Sulfite reductase ferredoxin-like | 243 | 572 |
| PF00118.27 | Cpn60_TCP1 | TCP-1/cpn60 chaperonin family | 1045 | 3024 |
| PF16732.8 | ComP_DUS | Type IV minor pilin ComP, DNA uptake sequence receptor | 78 | 197 |
| PF14339.9 | DUF4394 | Domain of unknown function | 37 | 49 |
| PF17210.6 | SdrD_B | SdrD B-like domain | 226 | 608 |
| PF04358.16 | DsrC | DsrC like protein - dissimilatory sulfite reductase | 130 | 349 |
| PF07382.14 | HC2 | Histone H1-like nucleoprotein HC2 | 146 | 202 |
| PF03951.22 | Gln-synt_N | Glutamine synthetase, beta-Grasp domain | 204 | 613 |
| PF17802.4 | SpaA | Prealbumin-like fold domain | 127 | 340 |
| PF14905.9 | OMP_b-brl_3 | Outer membrane protein beta-barrel family | 119 | 372 |
| PF13501.9 | SoxY | Sulfur oxidation protein | 119 | 237 |
| PF00216.24 | Bac_DNA_binding | Bacterial DNA-binding protein | 312 | 1145 |
| PF16842.8 | RRM_occluded | Occluded RNA-recognition motif | 54 | 85 |
| PF14537.9 | Cytochrom_c3_2 | cytochrome c3 | 360 | 1103 |

| | | | | |
|---|---|---|---|---|
| PF01478.21 | Peptidase_A24 | Type IV leader peptidase family | 86 | 217 |
| PF06841.15 | Phage_T4_gp19 | T4-like virus tail tube protein gp19 | 57 | 112 |
| PF00669.23 | Flagellin_N | Bacterial flagellin N-terminal helical region | 483 | 1670 |
| PF17228.5 | SGP | Sulfur globule protein | 80 | 127 |
| PF00034.24 | Cytochrom_C | cytochrome C | 531 | 2091 |
| PF00076.25 | RRM_1 | RNA recognition motif | 187 | 527 |
| PF04896.15 | AmoC | Ammonia monooxygenase methane monooxygenase, subunit C | 65 | 68 |
| PF03953.20 | Tubulin_C | Tubulin | 62 | 65 |
| PF03863.16 | Phage_mat-A | Phage maturation protein | 73 | 176 |
| PF19572.2 | PorV | Type IX secretion system protein PorV | 63 | 130 |
| PF03063.23 | Prismane | Prismane/CO dehydrogenase family | 216 | 481 |
| PF00267.24 | Porin_1 | Gram negative porin | 444 | 755 |
| PF00073.23 | Rhv | picornavirus capsid protein | 58 | 120 |
| PF00909.24 | Ammonium_transp | Ammonium transporter family | 368 | 839 |
| PF02461.19 | AMO | Ammonia monooxygenase/methane monooxygenase, subunit C | 81 | 83 |
| PF03143.20 | GTP_EFTU_D3 | Elongation factor Tu C-terminal domain | 396 | 1657 |

| | | | | |
|---|---|---|---|---|
| PF02530.17 | Porin_2 | porin subfamily | 307 | 356 |
| PF13433.9 | Peripla_BP_5 | periplasmic binding protein | 398 | 1717 |
| PF06348.14 | DUF1059 | Domain of unknown function | 37 | 60 |
| PF15511.9 | CENP-T_C | Centromere kinetochore component CENP-T histone fold | 21 | 23 |
| PF02635.18 | DrsE | intracellular small protein involved in sulfur reduction | 247 | 665 |
| PF13585.9 | CHU_C | CHU_C Type IX secretion signal domain | 271 | 460 |
| PF01618.19 | MotA_ExbB | MotA/TolQ/ExbB proton channel family | 228 | 858 |
| PF19516.2 | DUF6049 | Domain of unknown function | 19 | 20 |
| PF00166.24 | Cpn10 | Chaperonin 10 Kd subunit | 462 | 1387 |
| PF13360.9 | PQQ_2 | Outer membrane protein assembly factor BamB | 223 | 711 |
| PF00998.26 | RdRP_3 | Viral RNA dependent RNA polymerase | 103 | 152 |
| PF13179.9 | DUF4006 | Domain of unknown function | 31 | 41 |
| PF10518.12 | TAT_signal | TAT (twin-arginine translocation) pathway signal sequence | 379 | 1065 |
| PF00700.24 | Flagellin_C | Bacterial flagellin C-terminal helical region | 455 | 1554 |
| PF05124.15 | S_layer_C | S-layer like family, outer domain | 33 | 38 |

| | | | | |
|---|---|---|---|---|
| PF16982.8 | Flp1_like | S-layer like family, outer domain | 65 | 73 |
| PF07589.14 | PEP-CTERM | PEP-CTERM motif | 1216 | 1510 |
| PF07642.14 | BBP2 | Putative beta-barrel porin-2, OmpL-like. bbp2 | 105 | 183 |
| PF13370.9 | Fer4_13 | 4Fe-4S single cluster domain of Ferredoxin | 121 | 335 |
